# Spatial organization of bacterial sphingolipid synthesis enzymes

**DOI:** 10.1101/2023.11.08.566215

**Authors:** Chioma G. Uchendu, Eric A. Klein

## Abstract

Sphingolipids are produced by nearly all eukaryotes where they play significant roles in cellular processes such as cell growth, division, programmed cell death, angiogenesis, and inflammation. While it was previously believed that sphingolipids were quite rare among bacteria, bioinformatic analysis of the recently identified bacterial sphingolipid synthesis genes suggests that these lipids are likely to be produced by a wide range of microbial species. The sphingolipid synthesis pathway consists of three critical enzymes. Serine palmitoyltransferase catalyzes the condensation of serine with palmitoyl-CoA (or palmitoyl-acyl carrier protein), ceramide synthase adds the second acyl chain, and a reductase reduces the ketone present on the long-chain base. While there is general agreement regarding the identity of these bacterial enzymes, the precise mechanism and order of chemical reactions for microbial sphingolipid synthesis is more ambiguous. Two mechanisms have been proposed. First, the synthesis pathway may follow the well characterized eukaryotic pathway in which the long-chain base is reduced prior to the addition of the second acyl chain. Alternatively, our previous work suggests that addition of the second acyl chain precedes the reduction of the long-chain base. To distinguish between these two models, we investigated the subcellular localization of these three key enzymes. We found that serine palmitoyltransferase and ceramide synthase are localized to the cytoplasm whereas the ceramide reductase is in the periplasmic space. This is consistent with our previously proposed model wherein the second acyl chain is added in the cytoplasm prior to export to the periplasm where the lipid molecule is reduced.

## Introduction

Sphingolipids play a critical role in eukaryotic cells where they are involved in membrane structure and function and serve as important signaling molecules (1). By contrast, prokaryotic sphingolipids were considered to be rare, with notable exceptions in Bacteroidetes where they play a role in modulating the host immune response (2) as well as Sphingomonads where they functionally replace lipopolysaccharide (LPS) (3). One reason that bacteria were thought to lack sphingolipids is that, apart from serine palmitoyltransferase (Spt), they do not encode homologues of the synthetic enzymes used in eukaryotes.

In a study of adaptation to phosphate limitation in *Caulobacter crescentus*, we found that this organism is also capable of producing sphingolipids (4). Using a set of genetic screens, we and others identified the key genes required for bacterial sphingolipid synthesis (5,6) (Figure 1). In addition to the conserved *spt* gene, we identified genes encoding a reductase (*cerR*) and an acyl-transferase (*bcerS*). Our biochemical data, along with bioinformatic analyses, led us to propose a synthetic pathway in which addition of the second acyl chain occurs prior to lipid reduction (Figure 1) (6), which is distinct from the well-characterized eukaryotic pathway. However, others in the field have proposed that these enzymes do, in fact, perform this synthesis in a pathway akin to eukaryotes (Figure 1) (5).

**Figure 1.**
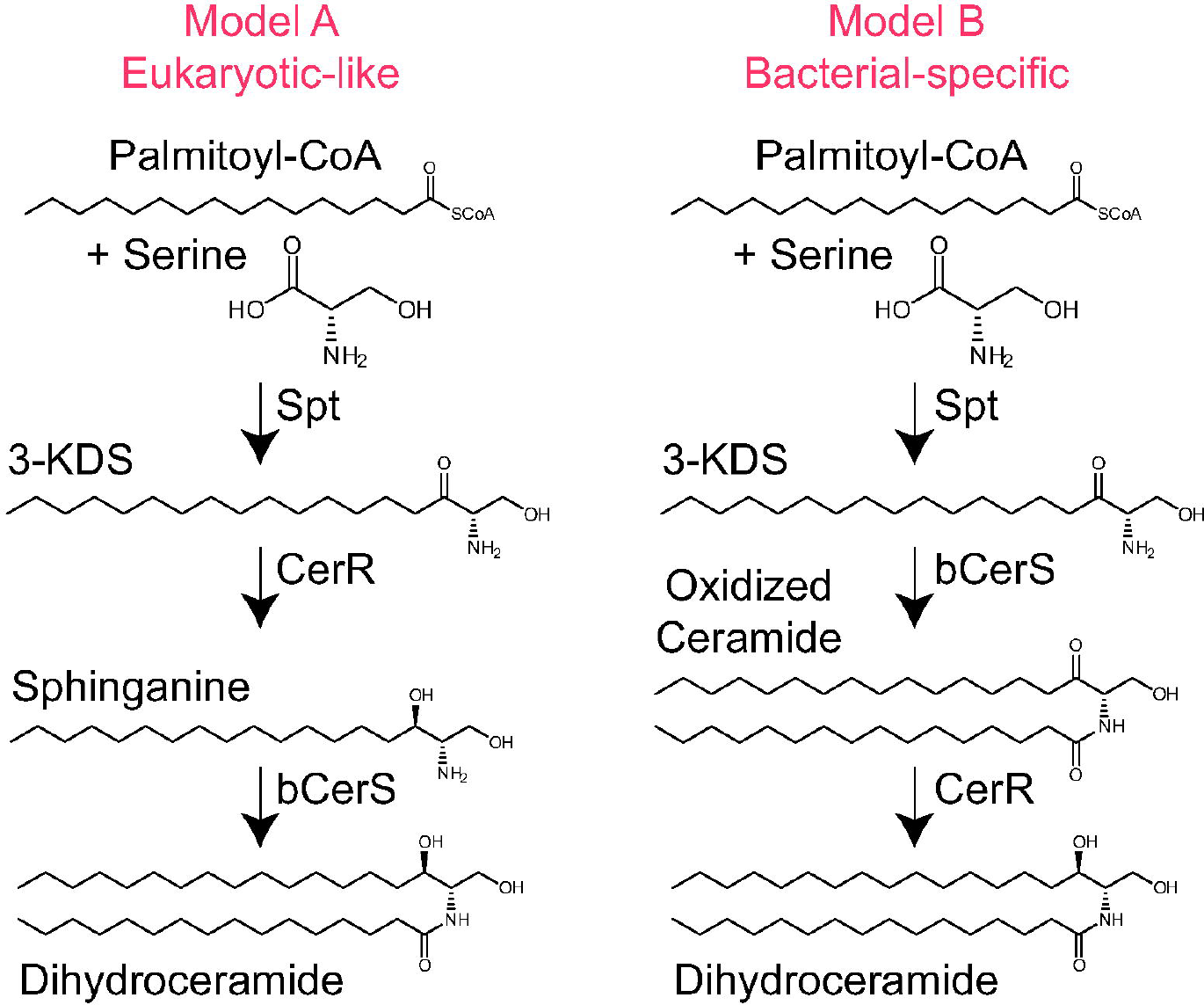
Proposed mechanisms of bacterial sphingolipid synthesis. Two recent studies identified the genes required for ceramide synthesis in *C. crescentus* (5,6). The proposed synthetic mechanisms either follow a similar chemistry to that found in eukaryotes (left) or operate in a bacterial-specific order (right).

To distinguish between these two proposed models, we have investigated the spatial organization of the sphingolipid synthesis enzymes. Using a set of orthogonal experimental approaches, we found that Spt and bCerS are cytoplasmic proteins, whereas CerR resides in the periplasm. Given that these lipids are ultimately trafficked to the outer membrane (7,8), our findings support the model in which Spt and bCerS act first in the cytoplasm prior to lipid translocation to the periplasm for subsequent reduction and trafficking to the outer membrane.

## Results

### Cell permeabilization reveals spatial arrangement of Sphingolipid synthesis enzymes

Mass spectrometry analysis of sphingolipid intermediates demonstrated the accumulation of oxidized-ceramide upon *cerR* deletion (6). The presence of a second acyl chain in this molecule suggests that either CerR acts after the addition of the second acyl chain, or bCerS acylates both 3-ketodihydrosphingosine (oxidized long-chain base) and sphinganine (reduced long-chain base). To distinguish between these two possibilities, we considered that these enzymes may occupy distinct subcellular niches. To assess protein localization, we repeated a previous experiment and expressed mCherry-tagged alleles of Spt, bCerS, and CerR and incubated the respective bacteria with chloroform-saturated Tris buffer, which preferentially permeabilizes the outer membrane and releases soluble periplasmic proteins (6,9). Spt and bCerS retained fluorescence upon permeabilization, indicating their potential localization in either the cytoplasm or inner membrane (Figure 2). By contrast, CerR showed a complete loss of signal, suggesting that it is most likely a soluble periplasmic protein (Figure 2). These results are consistent with our hypothesis that these enzymes are compartmentalized differently within the cell.

**Figure 2.**
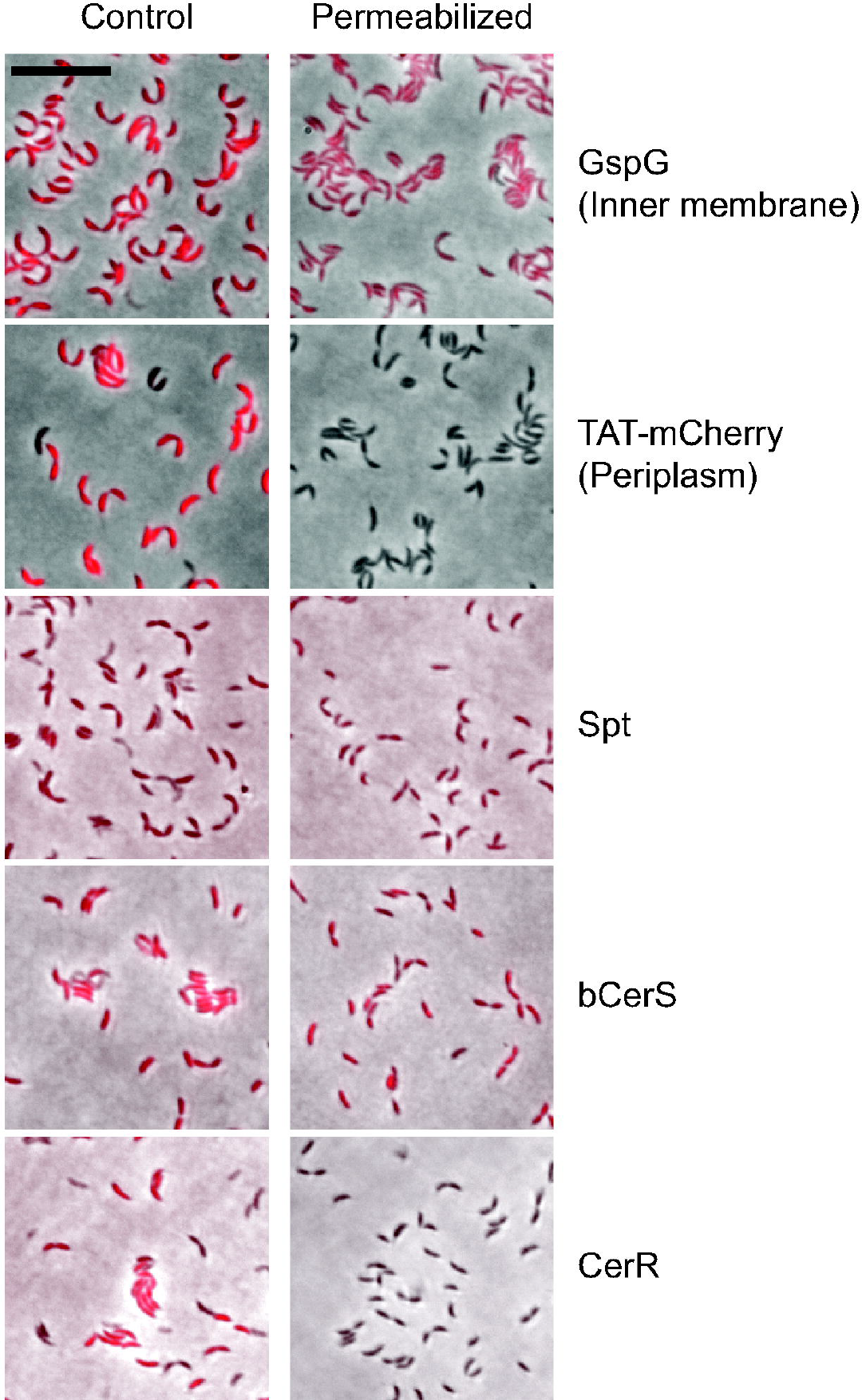
Permeabilization of the outer membrane suggests distinct subcellular localizations of the sphingolipid synthesis enzymes. Cells expressing the indicated mCherry-tagged proteins were grown overnight in the presence of 0.3% xylose or 0.5 mM vanillate to induce expression. GspG-mCherry (10) and TAT-mCherry (9) are control inner-membrane and periplasmic proteins, respectively. Control and permeabilized cells were visualized by fluorescence microscopy and the loss of fluorescence upon permeabilization was assessed. The images are the overlay of phase and fluorescent images. Scale bar = 10 μm.

### Beta-lactamase fusion assay to evaluate periplasmic protein localization

As an orthogonal approach to assess subcellular localization, we utilized beta-lactamase (bla) fusions as a probe for periplasmic secretion (10). The mechanism of action of beta-lactam antibiotics, such as carbenicillin, involves the inhibition of penicillin binding proteins (PBPs) in the periplasm of Gram-negative bacteria (11). To test for periplasmic localization, we made vanillate-inducible bla-fusions to the C-termini of Spt, CerR, and bCerS. We chose C-terminal fusions to avoid potentially disrupting any N-terminal signal sequence required for protein secretion. Additionally, *C. crescentus* is naturally resistant to carbenicillin due to the expression of beta-lactamase (12); therefore, we conducted all experiments using the *bla6* deletion strain (13) to restore beta-lactam sensitivity. Lastly, the fusion constructs lacked the *bla* signal sequence to ensure that any secretion was due solely to the sphingolipid-synthesis enzyme of interest. The respective strains were cultured both in the presence and absence of vanillate and/or carbenicillin. Growth of wild-type and Δ*bla6* strains on carbenicillin plates confirmed their respective antibiotic sensitivities (Figure 3). In the presence of both carbenicillin and vanillate, only the CerR fusion displayed growth (Figure 3), whereas Spt and bCerS fusions remained sensitive to carbenicillin (Figures 3). By contrast, expression of the FLAG-tagged enzymes did not restore carbenicillin resistance. These results are consistent with the data obtained by permeabilizing the outer-membrane (Figure 2) and support the periplasmic localization of CerR.

**Figure 3.**
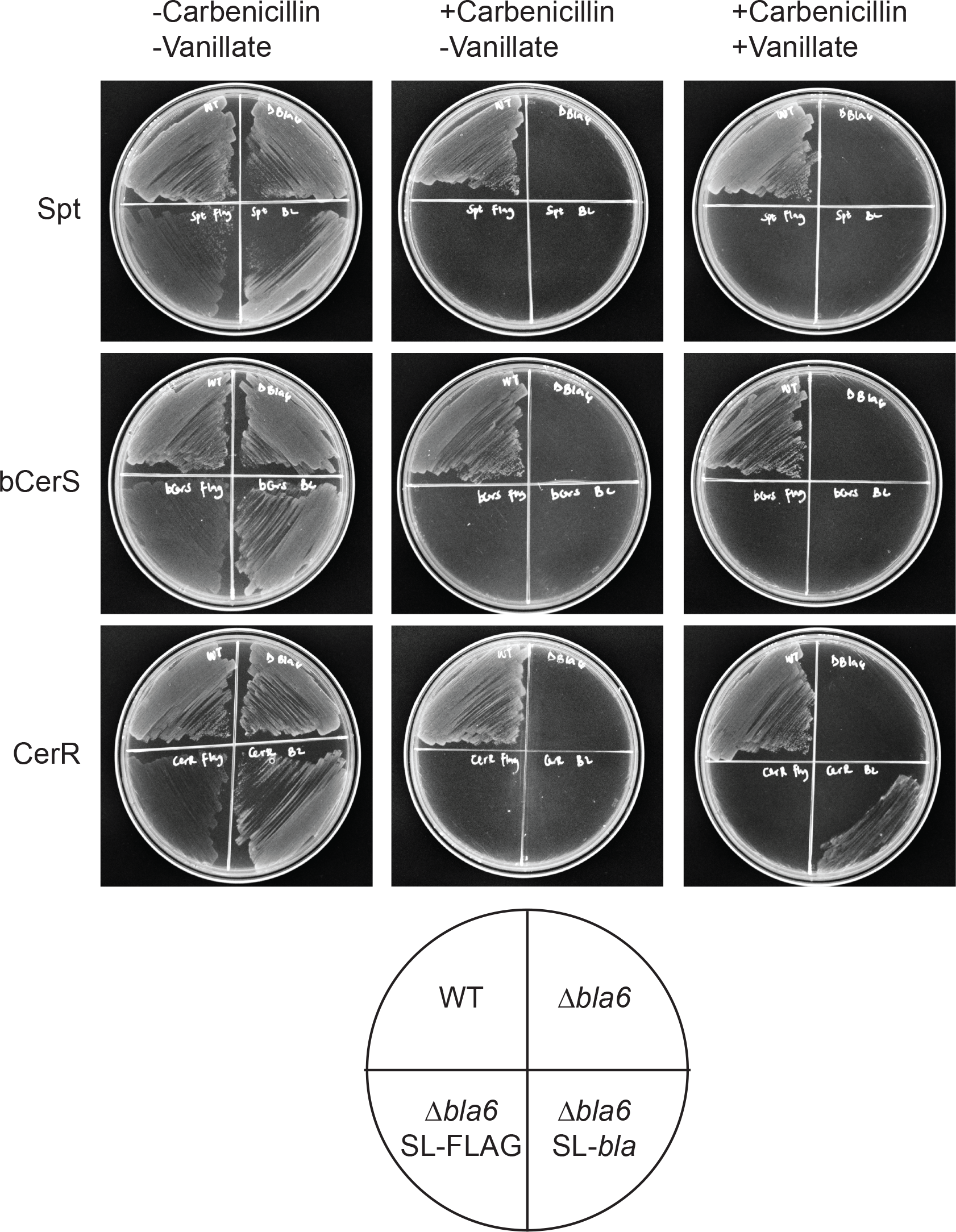
Beta-lactam resistance indicates periplasmic localization. The indicated strains were grown overnight +/- 1 mM vanillate to induce beta-lactamase fusion expression. Wild-type (carbenicillin resistant) and Δ*bla6* (carbenicillin sensitive cells) were included as controls. The strains were streaked onto PYE agar plates as diagramed and growth was assessed after 48 hr. Plates were either plain PYE (all strains should grow), PYE + carbenicillin (only WT should grow), and PYE + carbenicillin + vanillate (only WT and periplasmic beta-lactamase fusions should grow).

### Solubility of the sphingolipid synthesis proteins

The eukaryotic Spt, KDSR, and CerS enzymes are all intrinsic membrane proteins (14). Previous characterization of the bacterial Spt demonstrated that, by contrast, this enzyme is soluble (15). To determine the solubility of CerR and bCerS, we performed subcellular fractionation assays to separate soluble and membrane-associated proteins. Our analysis confirmed that Spt is soluble (Figure 4A). By contrast, both bCerS and CerR were found in both the soluble and membrane fractions (Figure 4B). Analysis of these protein sequences with CCTOP did not identify any predicted transmembrane regions (16); therefore, we questioned whether these proteins are integral or peripheral membrane proteins. To distinguish between these possibilities, we washed the membrane pellet with an alkaline sodium carbonate solution (0.1M Na2CO3, pH 11.0) which removes peripheral membrane proteins (17). Following this wash, both CerR and bCerS were found in the soluble fractions (Figure 4B), consistent with peripheral membrane association. Together with the localization data, we propose a model in which Spt is a soluble cytoplasmic protein, bCerS is a cytoplasmic peripheral membrane protein, and CerR is a periplasmic peripheral membrane protein (Figure 5).

**Figure 4.**
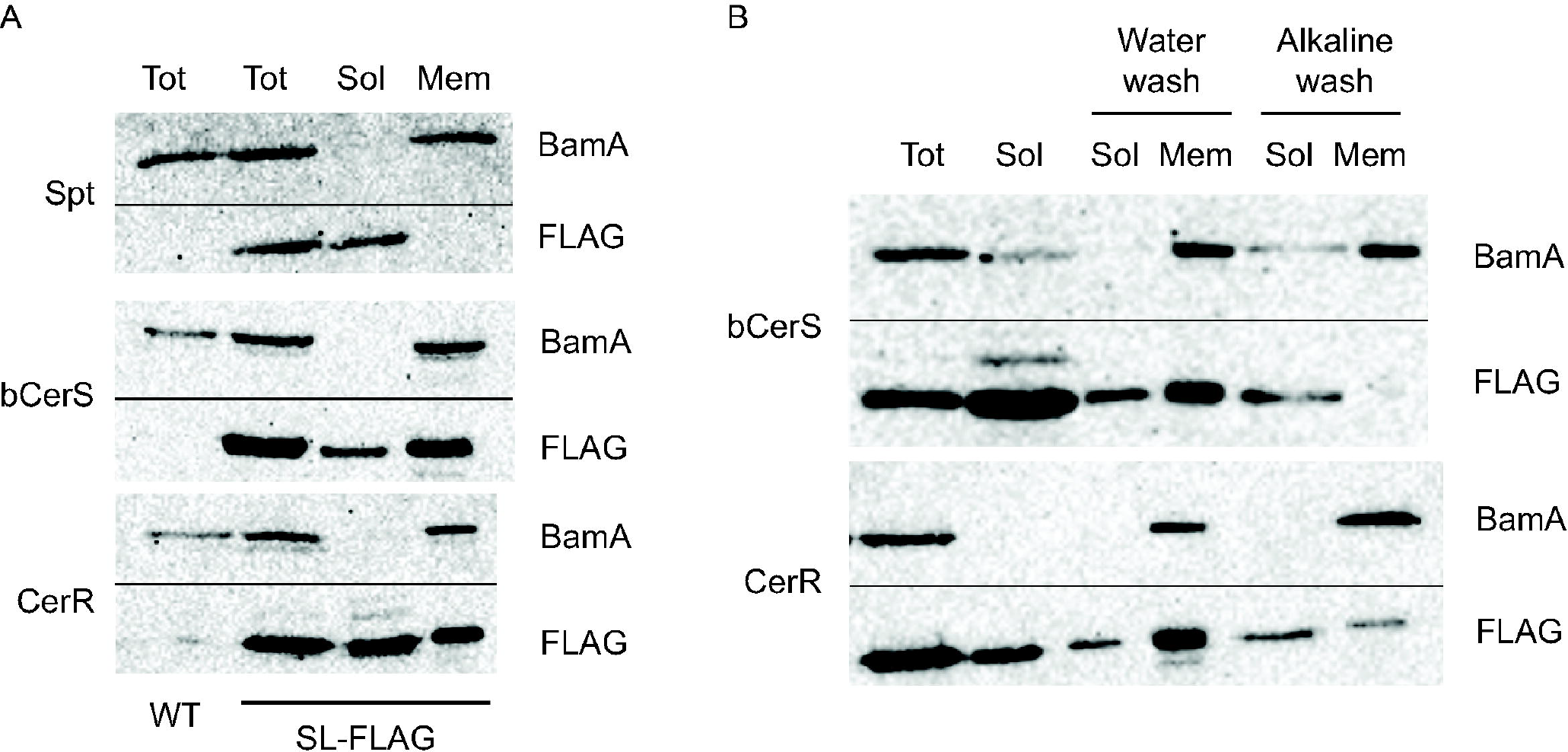
Subcellular fractionation identifies soluble and membrane-bound proteins. Cells expressing FLAG-tagged sphingolipid synthesis enzymes were grown overnight with 0.5 mM vanillate to induce expression. (A) Cells were lysed via French press and membranes were collected via ultracentrifugation. Total cell lysates (Tot), soluble proteins (Sol), and membrane proteins (Mem) were resolved by SDS-PAGE and analyzed by Western blot. BamA served as a membrane control protein. Wild-type (WT) cells did not express a FLAG-tagged enzyme as serves as an anti-FLAG negative control. (B) Membrane pellets collected as above were washed with either water or an alkaline solution (0.1M Na_2_CO_3_, pH 11) to release peripheral membrane proteins. Samples were analyzed by Western blot as above. Horizontal black lines indicate where the membranes were cut for antibody incubation.

**Figure 5.**
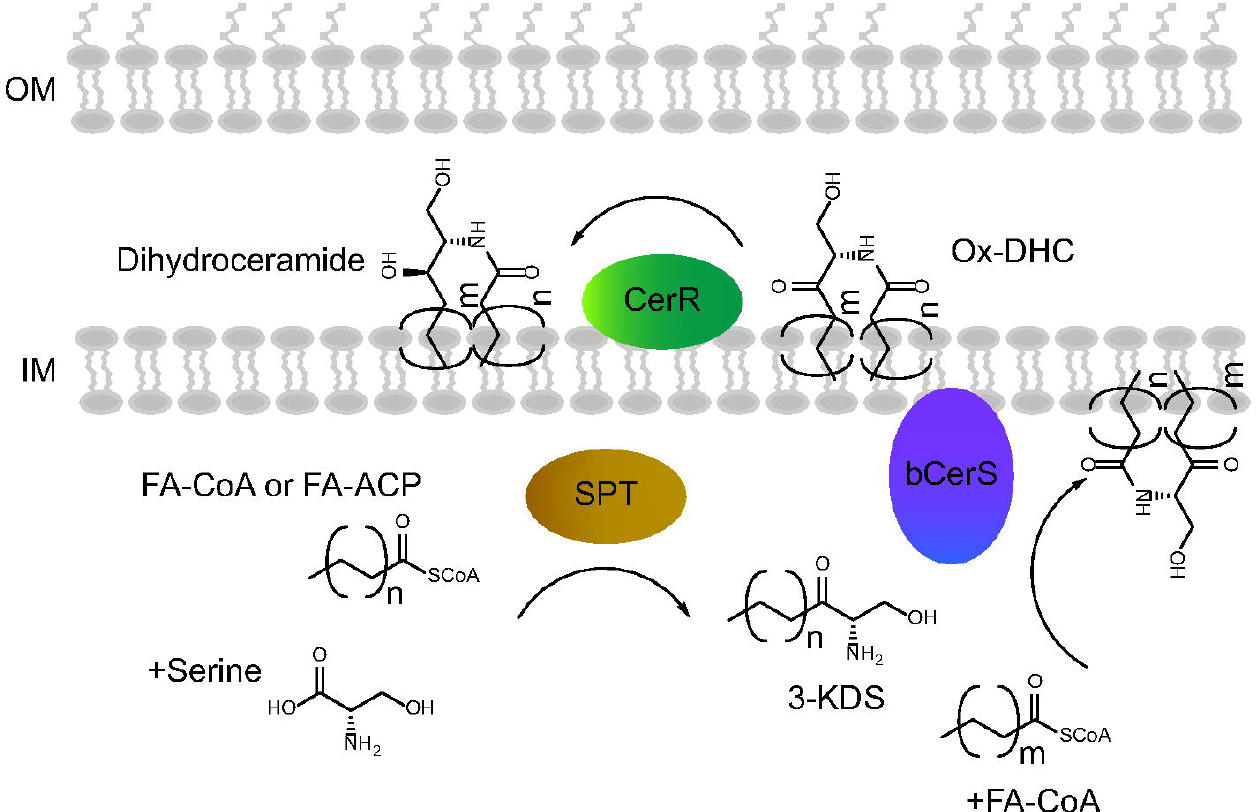
Model for bacterial sphingolipid synthesis. Based on our subcellular localization data, we propose that Spt is a soluble cytoplasmic enzyme that condenses either a fatty acid-CoA or a fatty acid-acyl carrier protein with serine to form 3-KDS. bCerS is a cytoplasmic peripheral inner-membrane protein that acylates 3-KDS to oxidized-ceramide/dihydroceramide (DHC). Lastly, CerR is a periplasmic peripheral membrane protein that reduces the oxidized sphingolipid to ceramide/DHC. While the synthetic enzymes are well characterized, we have not yet identified any flippases or lipid transporters required to move the sphingolipids across the various membrane compartments.

## Discussion

Although bacterial sphingolipids were previously thought to be rare, the recent elucidation of the microbial sphingolipid synthesis pathway suggests that these lipids may be present in a wide range of Gram-negative bacteria and smaller group of Gram-positive organisms (6). The physiological functions of these lipids appear to be species-specific. For example, *Bacteroides thetaiotaomicron* and *Porphyromonas gingivalis* use sphingolipids to suppress host inflammatory processes (2,18), *Sphingomonas* species replace lipopolysaccharide with glycosphingolipids (7), and *C. crescentus* sphingolipids protect against phage infection (4).

Further study of the physiological roles of bacterial sphingolipids will require a deeper understanding of the biochemical processes involved in their synthesis. Despite the discovery of the genes required for microbial sphingolipid production (5,6), there is disagreement regarding the order and nature of the biochemical reactions. One model suggests that the synthesis follows the same pathway as that of eukaryotes (5), where reduction of 3-KDS precedes addition of a second acyl-chain. Our model proposed that the second acyl-chain is added prior to reduction of the ceramide molecule (6). The data presented in this study aim to distinguish between these models. Based on independent assays of subcellular localization, we have demonstrated that Spt and bCerS are cytoplasmic while CerR is periplasmic (Figures 2-3). This spatial organization is consistent with our model that acylation occurs before lipid reduction. The alternative model would require that 3-KDS (the product of Spt) be translocated to the periplasm for reduction, the resulting sphinganine then be flipped back to the cytoplasm for acylation, and the final lipid being flipped one more time for final translocation to the outer membrane. This back-and-forth process for sphingolipid synthesis seems unlikely and is inconsistent with our previous finding that deletion of *cerR* results in the accumulation of an oxidized-ceramide metabolic intermediate (6).

Although multiple lines of evidence place CerR in the periplasm, this introduces a new question regarding the nature of the reducing equivalent. CerR has homology to the NDUFA9 protein, which is a component of mitochondrial Complex I and functions as an NADH-dependent ubiquinone reductase. The NADH binding site is conserved in CerR (6); however, there is no free NADH in the periplasm to catalyze this reaction. Given its peripheral membrane association (Figure 4B) and its homology to NDUFA9, we hypothesize that CerR may interact with either Complex I or another, yet unidentified, protein which enables electron transfer from NADH to oxidized-ceramide. A similar mechanism is used by the periplasmic nitrate reduction enzyme NarG, which receives electrons from NarI, an inner-membrane cytochrome *b* quinol oxidase (19). Further work will be required to identify CerR-interacting proteins.

While these core synthetic enzymes are found in all bacterial species that make sphingolipids, a potential 3-KDS reductase (KDSR) has been identified in *B. thetaiotaomicron* (20). This enzyme (BT_0972) can reduce 3-KDS to sphinganine *in vitro* and overexpression of BT_0972 converts 3-KDS to sphinganine when expressed in *E. coli*. These results suggest that, in this organism, there may be an additional sphingolipid synthesis pathway that runs parallel to that in eukaryotes. Whether this gene is involved in sphingolipid synthesis *in vivo* is not clear since it appears to be essential, and a loss of function mutant was unavailable for these experiments. As we learn more about the physiology of bacterial sphingolipids, we expect there to be many species-specific lipid metabolic pathways which contribute to the rich diversity of microbial sphingolipid molecules.

## Methods

### Bacterial strains, plasmids, and growth conditions

The specific strains, plasmids and primers utilized in this study can be found in Supplementary Tables 1-3. More information on the strain construction is also available in the supplementary materials. For routine culturing of *C. crescentus* wild type strain NA1000 and its derivatives, bacteria were cultured at 30 °C in peptone-yeast-extract (PYE) medium. *E. coli* strains were grown at 37 °C in Luria-Bertani (LB) medium. Selection antibiotics were added as required at the following concentration: 50 μg ml^-1^ of spectinomycin in broth and agar for *E. coli* and 25 μg ml^-1^ in broth and 100 μg ml^-1^ in agar for *C. crescentus*. Gene expression was induced in *C. crescentus* by the addition of 0.5 mM vanillate or 0.3% xylose.

### Outer membrane permeabilization and imaging

Chloroform-saturated Tris buffer was prepared by mixing 50 mM Tris, pH 7.4 with chloroform (70:30) and shaking the mixture at room temperature for 30 min. mCherry-fusion strains were induced with xylose or vanillate overnight, collected via centrifugation (2 min at 6,000 x *g*, 4 °C), and resuspended in an equal volume of the aqueous phase of the chloroform-saturated Tris buffer. Resuspended cells were rocked for 45 min at room temperature and then washed twice in 50 mM Tris, pH 7.4 (via centrifugation for 10 min at 5,000 x *g*) to remove residual chloroform.

Control cells were treated as above, but incubated in 50 mM Tris, pH 7.4 without chloroform. The permeabilized cells were spotted onto 1% agarose pads for imaging. Fluorescence microscopy was performed on a Nikon Ti-E inverted microscope equipped with a Prior Lumen 220PRO illumination system, CFI Plan Apochromat 100X oil immersion objective (NA 1.45, WD 0.13 mm), Zyla sCMOS 5.5-megapixel camera (Andor), and NIS Elements v, 4.20.01 for image acquisition.

### Beta-lactam resistance assay

The indicated strains were grown +/-vanillate overnight to induce expression of the beta-lactamase fusion proteins. Samples of the overnight cultures were streaked onto PYE agar plates containing +/-1 mM vanillate and +/-50 μg ml^-1^ carbenicillin. Bacterial growth was assessed after 48 hours of growth at 30 °C.

### Subcellular fractionation

Strains encoding FLAG-tagged alleles of the sphingolipid synthesis genes were grown in 500 ml PYE overnight with vanillate (0.5 mM). A 1 ml sample of the culture was removed as a total protein sample. The remainder of the culture was centrifuged at 5,000 x g for 10 minutes, the supernatant was discarded, and the pellet was resuspended in 3 ml TE buffer (10 mM Tris, pH 8, 1 mM EDTA) and lysed by 2-3 passages through a French press (20,000 psi). The lysed cells were centrifuged at 4 °C at 10,000 x g for 10 minutes to remove unbroken cells. The supernatant further centrifuged at 4 °C at 200,000 x g for 1 hour to pellet the membranes. Following protein concentration determination by BCA assay, the supernatant (soluble proteins), membranes, and total lysates were solubilized in 1X Laemmli buffer and denatured at 90 °C for 5 minutes.

### Removal of peripheral membrane proteins

Membrane pellets collected by ultracentrifugation were resuspended at a final concentration of 0.1 μg μl^-1^ with either water (control) or an alkaline solution (0.1M Na2CO3, pH 11) and incubated for 30 mins at 4 °C (17). The sample was centrifuged at 200,000 x g for 1 hour at 4 °C. After ultracentrifugation, the protein concentrations of the supernatant (peripheral membrane proteins) and pellet (intrinsic membrane proteins) were determined (BCA assay) and the samples were solubilized in 1X Laemmli buffer and denatured at 90 °C for 5 minutes.

### Western blotting

Proteins were resolved by SDS-PAGE on a 12% acrylamide gel with 20 μg of protein per well. After transferring proteins to a nitrocellulose membrane, target proteins were detecting using primary antibodies against the FLAG tag (proteintech; 20543-1-AP; 1:1000) and BamA (kind gift from Trevor Lithgow, Monash University; 1:50,000; (21)). Bands were detected using horseradish peroxidase conjugated secondary antibodies and ECL reagents (Cytiva) and imaged on a Bio-Rad Chemidoc MP.

## Data availability

All of the data for this work is contained within the manuscript.

## Acknowledgements

We thank Trevor Lithgow (Monash University, Australia) for providing the anti-BamA antibody.

## Author CrediT statement

**Chioma Uchendu:** Conceptualization, Methodology, Investigation, Writing-Original Draft, Visualization. **Eric Klein:** Conceptualization, Methodology, Investigation Writing-Original Draft, Visualization, Supervision.

## References

1. Harrison, P. J., Dunn, T. M., and Campopiano, D. J. (2018) Sphingolipid biosynthesis in man and microbes. Nat Prod Rep 35, 921–954

2. Brown, E. M., Ke, X., Hitchcock, D., Jeanfavre, S., Avila-Pacheco, J., Nakata, T., Arthur, T. D., Fornelos, N., Heim, C., Franzosa, E. A., Watson, N., Huttenhower, C., Haiser, H. J., Dillow, G., Graham, D. B., Finlay, B. B., Kostic, A. D., Porter, J. A., Vlamakis, H., Clish, C. B., and Xavier, R. J. (2019) Bacteroides-derived sphingolipids are critical for maintaining intestinal homeostasis and symbiosis. Cell Host Microbe 25, 668–680 e667

3. Kawahara, K., Moll, H., Knirel, Y. A., Seydel, U., and Zähringer, U. (2000) Structural analysis of two glycosphingolipids from the lipopolysaccharide-lacking bacterium Sphingomonas capsulata. European Journal of Biochemistry 267, 1837–1846

4. Stankeviciute, G., Guan, Z., Goldfine, H., and Klein, E. A. (2019) Caulobacter crescentus adapts to phosphate starvation by synthesizing anionic glycoglycerolipids and a novel glycosphingolipid. mBio 10, e00107–00119

5. Olea-Ozuna, R. J., Poggio, S., EdBergstrom Quiroz-Rocha, E., Garcia-Soriano, D. A., Sahonero-Canavesi, D. X., Padilla-Gomez, J., Martinez-Aguilar, L., Lopez-Lara, I. M., Thomas-Oates, J., and Geiger, O. (2021) Five structural genes required for ceramide synthesis in Caulobacter and for bacterial survival. Environ Microbiol 23, 143–159

6. Stankeviciute, G., Tang, P., Ashley, B., Chamberlain, J. D., Hansen, M. E. B., Coleman, A., D’Emilia, R., Fu, L., Mohan, E. C., Nguyen, H., Guan, Z., Campopiano, D. J., and Klein, E. A. (2022) Convergent evolution of bacterial ceramide synthesis. Nat Chem Biol 18, 305–312

7. Kawasaki, S., Moriguchi, R., Sekiya, K., Nakai, T., Ono, E., Kume, K., and Kawahara, K. (1994) The cell envelope structure of the lipopolysaccharide-lacking gram-negative bacterium Sphingomonas paucimobilis. J Bacteriol 176, 284–290

8. Zik, J. J., Yoon, S. H., Guan, Z., Stankeviciute Skidmore, G., Gudoor, R. R., Davies, K. M., Deutschbauer, A. M., Goodlett, D. R., Klein, E. A., and Ryan, K. R. (2022) Caulobacter lipid A is conditionally dispensable in the absence of fur and in the presence of anionic sphingolipids. Cell Rep 39, 110888

9. Stankeviciute, G., Miguel, A. V., Radkov, A., Chou, S., Huang, K. C., and Klein, E. A. (2019) Differential modes of crosslinking establish spatially distinct regions of peptidoglycan in Caulobacter crescentus. Mol Microbiol 111, 995–1008

10. Schlimpert, S., Klein, E. A., Briegel, A., Hughes, V., Kahnt, J., Bolte, K., Maier, U. G., Brun, Y. V., Jensen, G. J., Gitai, Z., and Thanbichler, M. (2012) General protein diffusion barriers create compartments within bacterial cells. Cell 151, 1270–1282

11. Lima, L. M., Silva, B., Barbosa, G., and Barreiro, E. J. (2020) Beta-lactam antibiotics: an overview from a medicinal chemistry perspective. Eur J Med Chem 208, 112829

12. Markiewicz, Z., Kuzma, M., and Kwiatkowski, Z. (1986) Mutants of Caulobacter crescentus resistant to beta-lactam antibiotics. Acta Microbiol Pol 35, 335–340

13. Evinger, M., and Agabian, N. (1977) Envelope-associated nucleoid from Caulobacter crescentus stalked and swarmer cells. J Bacteriol 132, 294–301

14. Gault, C. R., Obeid, L. M., and Hannun, Y. A. (2010) An overview of sphingolipid metabolism: from synthesis to breakdown. Adv Exp Med Biol 688, 1–23

15. Raman, M. C., Johnson, K. A., Yard, B. A., Lowther, J., Carter, L. G., Naismith, J. H., and Campopiano, D. J. (2009) The external aldimine form of serine palmitoyltransferase: structural, kinetic, and spectroscopic analysis of the wild-type enzyme and HSAN1 mutant mimics. J Biol Chem 284, 17328–17339

16. Dobson, L., Remenyi, I., and Tusnady, G. E. (2015) CCTOP: a Consensus Constrained TOPology prediction web server. Nucleic Acids Res 43, W408–412

17. Kongpracha, P., Wiriyasermkul, P., Isozumi, N., Moriyama, S., Kanai, Y., and Nagamori, S. (2022) Simple but efficacious enrichment of integral membrane proteins and their interactions for in-depth membrane proteomics. Mol Cell Proteomics 21, 100206

18. Moye, Z. D., Valiuskyte, K., Dewhirst, F. E., Nichols, F. C., and Davey, M. E. (2016) Synthesis of sphingolipids impacts survival of Porphyromonas gingivalis and the presentation of surface polysaccharides. Front Microbiol 7, 1919

19. Durand, S., and Guillier, M. (2021) Transcriptional and post-transcriptional control of the nitrate respiration in bacteria. Front Mol Biosci 8, 667758

20. Lee, M. T., Le, H. H., Besler, K. R., and Johnson, E. L. (2022) Identification and characterization of 3-ketosphinganine reductase activity encoded at the BT_0972 locus in Bacteroides thetaiotaomicron. J Lipid Res 63, 100236

21. Anwari, K., Poggio, S., Perry, A., Gatsos, X., Ramarathinam, S. H., Williamson, N. A., Noinaj, N., Buchanan, S., Gabriel, K., Purcell, A. W., Jacobs-Wagner, C., and Lithgow, T. (2010) A modular BAM complex in the outer membrane of the alpha-proteobacterium Caulobacter crescentus. PLoS One 5, e8619

