## Supplemental materials for "Spatial organization of bacterial sphingolipid synthesis enzymes"

**Supplementary Methods**

<u>Strain construction</u>

Strains for vanillate-inducible expression of FLAG-tagged sphingolipid synthesis genes were created by PCR amplifying *spt* (EK1306/1307), *bcerS* (EK1199/1200), and *cerR* (EK1158/1174) from NA1000 genomic DNA and ligating into the NdeI/NheI site of pVCFPC-1. The resulting plasmids were transformed into their respective gene knockout strains.

Strains for vanillate-inducible expression of b-lactamase fusions were created by PCR amplifying the following genes from NA1000 genomic DNA without the terminal stop codon: *spt* (EK1306/1312), *bcerS* (EK1199/1313), and *cerR* (1158/1309). The resulting DNA fragments were ligated into the NdeI/EcoRI site of pVCFPC-1. The beta-lactamase gene, without the first 69 bases encoding the signal sequence, was PCR amplified from pBAD18 (EK1311/1318) and ligated into the EcoRI/NheI site of the respective fusion plasmids. The resulting plasmids were transformed into the Δ*bla6* background.

**Supplementary Table 1.** Strains used in this study.

| Strain | Genotype | Construction | Source |
|---|---|---|---|
| *C. crescentus* | | | |
| NA1000 | Synchronizeable variant of wild-type *C. crescentus* strain CB15 | | (1) |
| GS32 | Δ*ccna_01220* (*spt*) | | (2) |
| GS140 | Δ*ccna_01222* (*cerR*) | | (3) |
| GS198 | Δ*ccna_01212* (*bcerS*) | | (3) |
| CU19 | Δ*ccna_01220*; P*vanA*:: *ccna_01220-FLAG* | Transformation of GS32 with pCU16 | This study |
| CU24 | Δ*ccna_01222*; P*vanA*:: *ccna_01222-FLAG* | Transformation of GS140 with pCU22 | This study |
| GS215 | Δ*ccna_01212*; P*vanA*:: *ccna_01212-FLAG* | | (3) |
| EK291 | P*xylX*:: *gspG-mCherry* | | (3) |
| EK308 | P*xylX*:: *TAT-mCherry* | | (3) |
| GS289 | P*vanA*:: *ccna_01220-mcherry* | | (3) |
| GS290 | P*vanA*:: *ccna_01222-mcherry* | | (3) |
| GS291 | P*vanA*:: *ccna_01212-mcherry* | | (3) |
| LS177 | Δ*bla6* | | MRK Alley (unpublished) |
| CU65 | *Δbla6* P*vanA*:: *ccna_01220-bla* | Transformation of LS177 with pGS281 | This study |
| CU70 | *Δbla6* P*vanA*:: *ccna_01222-bla* | Transformation of LS177 with pGS282 | This study |
| CU71 | *Δbla6* P*vanA*:: *ccna_01212-bla* | Transformation of LS177 with pGS283 | This study |
| CU75 | *Δbla6* P*vanA*:: *ccna_01220-FLAG* | Transformation of LS177 with pCU16 | This study |
| CU77 | *Δbla6* P*vanA*:: *ccna_01222-FLAG* | Transformation of LS177 with pCU22 | This study |
| CU76 | *Δbla6* P*vanA*:: *ccna_01212-FLAG* | Transformation of LS177 with pGS193 | This study |
| *E. coli* | | | |
| DH5a | Cloning strain | | Thermo Scientific |

**Supplementary Table 2.** Plasmids used in this study.

| Name | Description | Source |
|---|---|---|
| pVCFPC-1 | Vanillate-inducible expression, $Spect^R$ | (4) |
| pGS193 | pVCFPC-1-based plasmid for *ccna_01212-FLAG* expression, $Spect^R$ | (3) |
| pCU16 | pVCFPC-1-based plasmid for *ccna_01220-FLAG* expression, $Spect^R$ | This study |
| pCU22 | pVCFPC-1-based plasmid for *ccna_01222-FLAG* expression, $Spect^R$ | This study |
| pBAD18 | Source of beta-lactamase gene (*bla*) | (5) |
| pGS281 | pVCFPC-1-based plasmid for *ccna_01220-bla* expression, $Spect^R$ | This study |
| pGS282 | pVCFPC-1-based plasmid for *ccna_01222-bla* expression, $Spect^R$ | This study |
| pGS283 | pVCFPC-1-based plasmid for *ccna_01212-bla* expression, $Spect^R$ | This study |

**Supplementary Table 3.** Primers used in this study.

| Name | Sequence |
|---|---|
| EK1158 | tactcatATGGCGACTGACGCGCGC |
| EK1174 | tactgctagcTTActtgtcatcgtcatccttgtagtcGGCGACGATATTTTTGGGTAGCC |
| EK1199 | tactcatATGCCTTTCGACTCCACCAACG |
| EK1200 | tactgctagcTTActtgtcatcgtcatccttgtagtcCAGGGCCTTCTCGTAGACGC |
| EK1306 | tactcatATGGGGCTATTTGATAAGCACCTGG |
| EK1307 | tactgctagcTTActtgtcatcgtcatccttgtagtcGGCGCGGGCGCGCTTGAG |
| EK1309 | tactgaattcGGCGACGATATTTTTGGGTAGCC |
| EK1311 | tactgctagcctaCCAATGCTTAATCAGTGAG |
| EK1312 | tactgaattcGGCGCGGGCGCGCTTGAG |
| EK1313 | tactgaattcCAGGGCCTTCTCGTAGACGC |
| EK1318 | tactgaattcggtggaggtggatctCACCCAGAAACGCTGGTGAAAG |

**Supplementary references**


1. Nierman, W. C., Feldblyum, T. V., Laub, M. T., Paulsen, I. T., Nelson, K. E., Eisen, J. A., Heidelberg, J. F., Alley, M. R., Ohta, N., Maddock, J. R., Potocka, I., Nelson, W. C., Newton, A., Stephens, C., Phadke, N. D., Ely, B., DeBoy, R. T., Dodson, R. J., Durkin, A. S., Gwinn, M. L., Haft, D. H., Kolonay, J. F., Smit, J., Craven, M. B., Khouri, H., Shetty, J., Berry, K., Utterback, T., Tran, K., Wolf, A., Vamathevan, J., Ermolaeva, M., White, O., Salzberg, S. L., Venter, J. C., Shapiro, L., and Fraser, C. M. (2001) Complete genome sequence of *Caulobacter crescentus*. *Proc Natl Acad Sci U S A* **98**, 4136-4141
2. Stankeviciute, G., Guan, Z., Goldfine, H., and Klein, E. A. (2019) *Caulobacter crescentus* adapts to phosphate starvation by synthesizing anionic glycoglycerolipids and a novel glycosphingolipid. *mBio* **10**, e00107-00119
3. Stankeviciute, G., Tang, P., Ashley, B., Chamberlain, J. D., Hansen, M. E. B., Coleman, A., D'Emilia, R., Fu, L., Mohan, E. C., Nguyen, H., Guan, Z., Campopiano, D. J., and Klein, E. A. (2022) Convergent evolution of bacterial ceramide synthesis. *Nat Chem Biol* **18**, 305-312
4. Thanbichler, M., Iniesta, A. A., and Shapiro, L. (2007) A comprehensive set of plasmids for vanillate- and xylose-inducible gene expression in *Caulobacter crescentus*. *Nucleic Acids Res* **35**, e137
5. Guzman, L. M., Belin, D., Carson, M. J., and Beckwith, J. (1995) Tight regulation, modulation, and high-level expression by vectors containing the arabinose PBAD promoter. *J Bacteriol* **177**, 4121-4130
